# Route of oxytetracycline administration differentially impacts the growth and gut microbiome of pigs co-infected with *Bordetella bronchiseptica* and *Pasteurella multocida*

**DOI:** 10.1101/2022.04.18.488710

**Authors:** Kathy T. Mou, Julian Trachsel, Amali Stephens, Nicole Ricker, Susan L. Brockmeier, Heather K. Allen, Crystal L. Loving

**Affiliations:** Oak Ridge Institute for Science and Education, Oak Ridge, Tennessee, USA; Food Safety and Enteric Pathogens Research Unit, National Animal Disease Center, ARS, USDA, Ames, IA, USA; Department of Pathobiology, Ontario Veterinary College, University of Guelph, Guelph, Ontario, Canada.; Virus and Prion Diseases Research Unit, National Animal Disease Center, ARS, USDA, Ames, IA, USA

**Author notes:** Address correspondence to Crystal L. Loving. current address.

**Keywords:** *Bordetella bronchiseptica*, *Pasteurella multocida*, oxytetracycline, microbiome, antimicrobial resistance genes, swine

## Abstract

Along with judicious antibiotic use, there is great interest in how the dose regimen of an antibiotic affects the animal gut microbiota. This study evaluated the impact of experimental respiratory infection alone or respiratory infection followed by oxytetracycline (oxytet) treatment on the animal’s health and its fecal microbiome. Piglets of approximately three weeks-of-age were separated into four groups (*n=*20 per group). One group remained non-infected and administered non-medicated feed and the other three groups were infected with *Bordetella bronchiseptica* (day 0) and *Pasteurella multocida* (day 4), with one group receiving non-medicated feed and the remaining two groups receiving oxytetr starting on day 7 by injection or in-feed (day 7-14). Infection with *B. bronchiseptica* and *P. multocida* negatively impacted piglet growth and induced mild pneumonia. Infection alone had minimal effect on the fecal microbiota community. When oxytet was administered either by injection or in-feed to treat the respiratory infections, both routes had minimal effect on clearing *B. bronchiseptica* and *P. multocida* in the animal. However, both routes appeared to limit lung lesion severity, and injected oxytet reduced the negative impact of infection on weight gain. Both routes had limited impact on the animal’s overall gut microbiome, including relative abundances of bacterial taxa and antibiotic resistance genes *tet32, tetW,* and *aph2*. Overall, oxytet administered by either route did not clear the respiratory infection, but oxytet administration minimized the negative health impacts of infection and had minor impact on the pig gut microbiome.

**Importance:** Efforts to address antibiotic resistance calls for improved antibiotic stewardship, including considering antibiotic administration route. While our previous study found in-feed oxytet had greater impact on the gut microbiome of healthy piglets than injected oxytet, it remained unknown if oxytet treatments would have the same impact on the microbiota of infected piglets. We evaluated the impact of respiratory infection alone or respiratory infection followed by oxytet treatment on the animals’ health and their gut microbiome profile. Respiratory infection negatively affected piglets’ health, but infection alone had minimal impact on the gut community. When oxytet was administered either in-feed or by injection to treat the respiratory infection, neither route of administration led to the clearance of the respiratory pathogens. However, oxytet minimized the negative health impacts of infection, and had minor impact on the pig gut microbiome. These findings are informative for disease management in food animals while integrating antibiotic stewardship practices.

## Introduction

To respond to the continually growing global health crisis of antibiotic resistance, international agencies and individual countries developed comprehensive action plans and introduced regulations to address antibiotic resistance and emphasize antibiotic stewardship (1). Though the relative contribution of antibiotic use in food animals to the global antimicrobial resistance crisis is poorly defined, in the U.S. regulation on veterinary antibiotic usage has increased in the last several years. For instance, the Food and Drug Administration established the Veterinary Feed Directive stating that medically important antimicrobial drugs for food-producing animals are only approved for use for disease treatment and prevention (2).

One consideration of antibiotic stewardship is to treat animals with an antibiotic by a route that has limited disturbance to the gastrointestinal (GI) microbiome (3), but is still effective against the cause of disease. Distinct dosing regimens can have significant differential effects on the GI microbiome. In swine, oral antimicrobial drugs have substantial impact on the GI microbiota by shifting the microbiota towards increased abundance of resistant or opportunistic pathogens, contributing to the emergence and transmission of antimicrobial resistance genes, and altering animal health (4–6). In one study, piglets administered oral ampicillin excreted higher levels of bla_TEM_ genes in feces than piglets administered ampicillin by intramuscular injection (7). In another study, there was a drastic impact on the relative abundance of specific GI bacterial members and availability of metabolites in the gut of weaned piglets fed a diet high in analytical grade zinc oxide (ZnO) in contrast to pigs fed a diet low in ZnO (8). Prophylactic administration of in-feed oxytetracycline (oxytet) to nursery age pigs for 7 days had a much larger impact on the GI microbial community structure and led to higher abundance of antibiotic resistance genes in pig feces than a single intramuscular injection of oxytet (3). Collectively, these results demonstrate that the impacts of antimicrobials on the overall GI microbial community and on the abundance of resistance genes can be reduced when the antimicrobial is administered by intramuscular injection than by in-feed.

Post-weaned piglets in the nursery are highly susceptible to disease, and prophylactic antimicrobials, including antibiotics, are commonly administered to prevent disease in healthy animals (9–11). While prior studies suggest a differential impact on microbiota with different antibiotic administration routes, the studies were completed in healthy animals (3, 4, 7, 8).

Respiratory and gastrointestinal infections are also known to significantly alter the gut microbial community in pigs (12). When antibiotics are administered to treat a disease, it remains unclear if the dosing route would have the same impact on the microbiota of infected animals as compared to healthy animals.

Oxytet is a broad-spectrum antibiotic available for administration via the oral and injected routes. Though it does modulate the microbiota of otherwise healthy nursery pigs (3), it is unclear if the exact dosing regimen would have the same modulating effect if administered to infected animals. Oxytet is used within the U.S. swine industry for the therapeutic treatment of multiple diseases, including respiratory disease caused by *Pasteurella multocida* (13, 14). It is difficult to reproduce *P. multocida* disease without a predisposing agent, such as porcine reproductive and respiratory syndrome virus or *Bordetella bronchiseptica* (15). Thus, the present study inoculated post-weaned piglets with two respiratory bacterial pathogens, *B. bronchiseptica* followed by *P. multocida*, to accomplish the following objectives: (1) investigate how respiratory infection alone affects the GI microbial community (using feces as a proxy for the GI microbiome and, (2) determine which antibiotic administration route would limit *P. multocida* disease with minimal effect on the gut microbiome of the animal. We hypothesized infection with *B. bronchiseptica* and *P. multocida* would negatively affect the health of the animal and significantly alter the gut microbiome. In addition, we hypothesized that both oral and injected oxytet would effectively treat the infection and, similar to the previous study (3), injected oxytet would have less impact on the gut microbiome. Overall, the goal was to provide evidence for a dosing method (oral or in-feed) that would be effective against *P. multocida* with limited disturbance to the gut microbial community and relative abundance of antimicrobial resistance genes.

## Materials and Methods

### Bordetella bronchiseptica *and* Pasteurella multocida *inocula*

*B. bronchiseptica* strain KM22, a virulent phase I swine isolate (16, 17), was prepared and administered to pigs as previously described (18). *Pasteurella multocida* serovar A:3 strain 4317, a non-toxigenic strain isolated from the lung of a pig with pneumonia from Arkansas in 1980, was similarly prepared and administered to pigs as previously mentioned (17), administering approximately 10^8^ colony forming units (CFU/mL) per animal. Susceptibility testing of *B. bronchiseptica* strain KM22 and *P. multocida* serovar A:3 strain 4317 was conducted through the Iowa State University Veterinary Diagnostic Laboratory and the minimum inhibitory concentration of the two strains to oxytetracycline was 0.5 µg/mL.

### Experimental Design

Animal studies were conducted in accordance with the recommendations in the Guide for the Care and Use of Laboratory Animals after review and approval by USDA-National Animal Disease Center Animal Care and Use Committee. A total of eighty 18-21-day-old mixed breed pigs arrived on site from a high health status herd (free of influenza A virus and porcine reproductive and respiratory syndrome virus) and randomly distributed to four treatment groups (*n* = 20 pigs/group). The weights of pigs were recorded on day of arrival (Day -1). Two pens per treatment group were established in a BSL-2 containment room to evaluate the impact of pen effect on observed differences, of which there were none. The control group was administered phosphate-buffered saline (PBS) instead of *B. bronchiseptica* and *P. multocida*, and received non-medicated feed (non-infected/non-medicated, NONINFnm). The three other groups consisted of pigs that were first infected with *B. bronchiseptica* strain KM22 (Day 0) and then *P. multocida* strain 4317 four days later (Day 4). On day 7, infected (INF) animals were then separated into three groups. One INF group was given feed containing oxytet (Terramycin® 200, Phibro, Teaneck, NJ) at the therapeutic dose of 10 mg/lb of body weight (assumed 17 lb pig and 450 g feed per pig per day) in a single trough per pen (INFfeed). The second INF group was administered a single intramuscular injection of oxytet (Liquamycin® LA-200 ®, Zoetis, Parsippany, NJ) at the recommended dose of 9 mg/lb based on the estimated weight of 17 lb to calculate injected dose (INFinject). The INFinject group received non-medicated feed for the duration of the study. The third INF group was not medicated and received non-medicated feed (INFnm).

On days 7, 11 and 14 (Figure 1), antemortem samples were collected from all pigs, including pig weight, nasal wash, plasma, and feces (exception, weights were not collected on day 7). Nasal wash was first collected as described (19) followed by a nasal swab. The nasal wash effluent and nasal swab were placed in a 15 mL conical tube, transported on ice, and aliquoted for downstream analysis of oxytet concentration level and bacterial enumeration.

Plasma was collected as previously described (20) into an EDTA blood tube. Feces or rectal swabs were collected from all pigs on days 7, 11, and 14, transported on ice to the lab, aliquoted for downstream applications, and stored at -80°C, as described previously (20). Fecal DNA was extracted from 195 samples using the PowerMag Microbiome DNA/RNA Kit (MoBio Laboratories, Carlsbad, CA).

On day 11, ten pigs from each group (5 pigs from each pen) were euthanized with a lethal dose of pentobarbital and necropsies were performed. On day 14, the remaining pigs were euthanized. Postmortem samples included tonsil tissue, tracheal wash, lung lavage, and lung tissue. Gross lung lesion scores were determined based on the percentage of each lung lobe affected and the percentage of total lung volume each lobe represented, calculated as described previously (21). Approximately 10 g of right middle lung lobe was placed in a ziploc bag and stored at -80 °C for assessing oxytet levels described below. Tonsil and lung tissue samples were dissected to approximately 1 g pieces and placed into gentleMACS C Tubes (Miltenyi Biotec, Auburn, CA). The weight of each sample was recorded and PBS was added to a 10% weight/volume. Samples were then homogenized using the gentleMACS Octo Dissociator (Miltenyi Biotec, Auburn, CA) for bacterial enumeration. Lung lavage and tracheal wash samples were collected using 50 mL and 3 mL sterile 1X PBS, respectively, following methods previously described (19). Lung lavage and tracheal wash were aliquoted for bacterial enumeration. Taking the weight data measured on days 0, 11, and 14 of the study, the average daily gain (ADG) of each pig was calculated over 11 and 14 days by (Weight on Day X – Weight on Day 0) / (Day X + 1). Lung lesion scores and average daily gain (ADG) were assumed parametric and analyzed using one-way analysis of variance (ANOVA) with Tukey’s multiple comparisons test in GraphPad Prism (La Jolla, CA, USA) to determine whether there were significant differences between the four treatment groups.

### B. bronchiseptica and P. multocida enumeration

The number of colony forming units (CFU) of *B. bronchiseptica* or *P. multocida* per mL of nasal wash, tonsil tissue homogenate, lung tissue homogenate, lung lavage, and tracheal wash were determined as previously described (19). Briefly, tracheal and lung samples were serially diluted in PBS and plated on blood agar. Nasal and tonsil samples were serially diluted in PBS and plated on blood agar supplemented with 2 µg/mL amikacin, 4 µg/mL vancomycin, and 4 µg/mL amphotericin B. The limit of detection was log_10_ of 1 (CFU of 10). Aliquots of tracheal and lung samples were also plated on brain-heart infusion agar supplemented with 0.01% NAD (w/v) and 5% horse serum to rule out aerobic bacterial infection (other than *B. bronchiseptica* and *P. multocida* in respective groups) to confirm lack of co-infection and no growth from other organisms was detected. To assess if there were differences in *B. bronchiseptica* or *P. multocida* colonization as a result of the route of oxytet treatment given (between INFnm, INFfeed, and INFinject), CFU/mL of each challenge isolate from antemortem samples on day 7 (taken before antibiotic administration), and postmortem samples on days 11 and 14 was log transformed and statistically analyzed using one-way ANOVA and Tukey’s multiple comparisons test in GraphPhad Prism (La Jolla, CA, USA). A *p*-value of 0.05 or less was considered significant.

### Oxytet concentrations

Concentrations of oxytet in porcine plasma, lung tissue, and nasal wash were determined using liquid chromatography mass spectrometry (LC-MS) at the Iowa State University Veterinary Diagnostic Laboratory Analytical Chemistry Services building on methods described previously (3). LC-MS/MS analysis for porcine plasma was performed using the Q Exactive Focus Hybrid Quadrupole-Orbitrap Mass Spectrometer coupled to a Dionex Ultimate 3000 (Thermo Fisher Scientific, San Jose, CA, USA). For lung tissue samples, LC-MS/MS analysis was performed using the TSQ Altis Triple Quadrupole Mass Spectrometer (Thermo Fisher Scientific, San Jose, CA) coupled to a Vanquish Flex UHPLC^+^ system consisting of a binary pump, autosampler, and column heater (Thermo Fisher Scientific, San Jose, CA). LC-MS/MS analysis for nasal wash samples were performed using the Finnigan TSQ Quantum Discovery MAX Triple Quadrupole Mass Spectrometer with an electrospray interface, a Finnigan Surveyor MS Pump Plus HPLC pump, and a Finnigan Surveyor Autosampler Plus autosampler (Thermo Fisher Scientific, San Jose, CA).

Plasma samples and the respective standards and quality control (QC) samples were prepared as described previously (3). To prepare lung tissue samples, standards, and QC samples for analysis, two grams of lung tissue from each animal were used. Demeclocycline was added at a concentration of 50 ng/g, along with 10 mL acetonitrile:water (4:1). Samples were vortexed for 5 minutes and centrifuged at 3000 rcf for 5 minutes. Supernatant was transferred to a tube containing 0.5 g of C18 silica and then 10 mL hexane was added. Samples were vortexed and centrifuged at 3000 rcf for 5 min again. The hexane was discarded and remaining supernatant was transferred into a test tube where the solvent was evaporated to dryness at room temperature. Samples were reconstituted with 0.075 mL 25% acetonitrile and 0.075 mL water and transferred to autosampler vials. Samples were centrifuged at 2500 rpm for 20 min prior to LC-MS analysis. Nasal wash samples were prepared by pipetting 0.1 mL of sample into a 1.5 mL microcentrifuge tube with 0.005 mL of 20000 ng/mL demeclocycline in 12.5% acetonitrile/0.1% formic acid. Each sample was vortexed and centrifuged at 2400 rpm for 20 min prior to LC-MS analysis.

For LC-MS analysis of porcine plasma, the injection volume was set to 0.002 mL. The mobile phases consisted of A: 0.1% formic acid in water and B: 0.1% formic acid in methanol at a flow rate of 0.3 mL/min. The mobile phase began at 7.5% B with a linear gradient to 95% B at 0.5 min - 6.5 min, followed by re-equilibration to 7.5% B. Separation was achieved with an Accucore C18 column, 100 mm x 2.1 mm, 2.6 µm particles (Thermo Scientific, San Jose, CA, USA) maintained at 35 °C. The retention time for oxytet and demeclocycline were 4.2 and 4.5 min, respectively. Full scan MS with wideband activation was used for analyte detection and three fragment ions were used for quantitation of each analyte species. The fragment ions for oxytet were at 461.155, 426.120, and 443.150 m/z, while fragment ions for demeclocycline were at 465.106, 443.950, and 448.080 m/z. Sequences consisting of porcine plasma blanks, calibration spikes, QC’s, and porcine plasma samples were batch processed with the same processing method in the Xcalibur software (Thermo Scientific, San Jose, CA) as previously described (3). Nine calibration spikes were prepared in blank porcine plasma covering the concentration range of 5 to 2,000 ng/mL oxytet. Calibration curves exhibited a correlation coefficient (r^2^) that exceeded 0.99. QC samples at 25, 75, and 750 ng/mL were within tolerance of ±15% of the nominal value. The limit of quantitation (LOQ) of the analysis was 10 ng/mL.

For LC-MS analysis of lung tissue, the injection volume was set to 0.002 mL. The mobile phases consisted of A: 0.05% formic acid in water and B: 0.05% formic acid in acetonitrile:methanol:water (45:45:10) at a flow rate of 0.3 mL/min. The elution gradient was held at 2% B for the first 0.5 min, 2 – 20% B from 0.5 to 1.2 min, 20 – 100% B from 1.2 to 6 min, held at 100% B from 6 to 6.5 min, and 100 – 2% B from 6.5 to 7.5 min, and held at 2% B for 0.5 min. Chromatographic separation was carried out on an Accucore^TM^ RP-MS, 100 mm × 2.1 mm, 2.6 µm particles (Thermo Fisher Scientific, San Jose, CA, USA), without a guard column, and maintained at 40^0^C. The retention time for oxytet and demeclocycline were 2.97 and 3.29 min, respectively. The fragment ions for identification and quantification of oxytet were at 461.155, 426.08, and 443.155 m/z, while fragment ions for demeclocycline were at 465.2, 430.05, and 448.08 m/z. Sequences consisting of lung tissue blanks, calibration spikes, QC’s, and lung tissue samples were batch processed with Xcalibur software (Thermo Scientific, San Jose, CA). A calibration curve was calculated using Quan Browser and a weighted (1/x) linear regression analysis was used to determine the slope, intercept, and correlation coefficient (r^2^).

Sample data were only processed if the value of r^2^ was > 0.999 and the calibrators and QCs (at 15, 25, and 150 ng/g) used were within a tolerance of ±15% of the nominal value. Nine calibration spikes were prepared in blank lung tissue covering the concentration range of 5 to 2,000 ng/mL oxytet. The LOQ was based on the calibration curve and was set at 5 ng/mL. For LC-MS analysis of nasal wash, the injection volume was set to 0.025 mL. The mobile phase consisted of A: 0.1% formic acid in water and B: 0.1% formic acid in acetonitrile at flow rates ranging from 0.250 to 0.350 mL/min. The mobile phase composition was initially 92.5% A and 7.5% B. After 8.5 min, the composition was changed to 95 % B. At 11.5 min, the gradient returned the composition to 7.5% B. Separation was accomplished using a Kinetex XB-C18 column, 100 mm x 2.1 mm, 2.6 µm (Phenomenex, Torrance, CA) and maintained at 45.0°C.

Retention times were 4.30 min for oxytet and 4.84 min for demeclocycline. The fragment ions for identification and quantification of oxytet were at 201, 226, and 426 m/z, while fragment ions for demeclocycline were at 154, 289, and 448 m/z. Instrumentation was controlled and data were handled using Xcalibur (Thermo Scientific, San Jose, CA). Fourteen calibration spikes were prepared in blank porcine plasma covering the concentration range of 0 to 800 ng/mL oxytet. The LOQ of the analysis was 10 ng/mL.

Oxytet concentrations from plasma, lung tissue, and nasal wash samples on days 11 and 14 between the four treatment groups were assessed using one-way ANOVA with Tukey’s multiple comparisons test. Oxytet concentrations from the same tissues between days 11 and 14 within each group were assessed using the Mann-Whitney test. All statistical calculations were done using GraphPad Prism (La Jolla, CA, USA), where a *p*-value of 0.05 or less was considered significant.

### Fecal microbiota sequencing and statistical analysis

The V4 region of the 16S rRNA gene was amplified from extracted fecal DNA and sequenced on a MiSeq (Illumina, San Diego, CA) using 2 x 250 V2 chemistry following the MiSeq Wet Lab SOP [(22); https://github.com/SchlossLab/MiSeq_WetLab_SOP/blob/master/MiSeq_WetLab_SOP_v4.md]. 16S rRNA raw fastq data were clustered into operational taxonomic units (OTUs) with ≥ 97% similarity in mothur using the MiSeq SOP protocol [(22); https://www.thur.org/wiki/MiSeq_SOP, accessed January 2020], with the addition of removing singletons and doubletons using the split.abund command (cutoff = 2). 16S rRNA gene sequences were aligned to the SILVA reference alignment. Data analysis and figure generation were done using R version 4.0.2, including the packages vegan, phyloseq, philentropy, and tidyverse for alpha- and beta-diversity measures of the samples from each treatment group (NONINFnm, INFnm, INFfeed, INFinject). Select groups were analyzed together to address two sets of research questions: 1) comparing NONINFnm and INFnm 16S sequence data to assess how co-infection of pigs with *B. bronchiseptica* and *P. multocida* impact gut microbial communities (reflected in feces), and 2) comparing INFnm, INFfeed, and INFinject 16S sequence data to assess how the oxytet dosing regimen used for treating *B. bronchiseptica* and *P. multocida* co-infection affected fecal microbial communities. Alpha-diversity metrics were calculated as previously described (23). The effects of oxytet treatment, and *B. bronchiseptica* and *P. multocida* colonization on the fecal microbial community structure within each sampled day were analyzed by calculating Bray-Curtis dissimilarities on rarefied OTU tables (1300 sequences per sample). Non-metric multidimensional scaling (NMDS) ordination based on Bray- Curtis dissimilarities, PERMANOVA pairwise comparisons of treatment groups using the pairwise.adonis function (R vegan package) (24) and p.adjust function (R stats package) were conducted as previously performed (23) to identify any significant differences in bacterial composition between groups on a given day. Significance was set at *P* < 0.05. To identify differentially abundant taxa between groups on a given day, the DESeq2 package (25) was used. Prior to testing, OTUs with fewer than 10 counts globally and samples with fewer than 1300 sequences were removed and the remaining OTUs agglomerated at the order level. The resulting counts were used as the input for DESeq2 in accordance with the package recommendations.

Shrinkage of log2-fold change estimates were applied using the apeglm method (26). OTUs were further classified at the order level. The significance of a log2-fold change value was determined using Wald test with parametric fit, and *p*-values ≤ 0.05 were considered significant. The null hypothesis was that relative abundance of taxon was not different between treatment groups on a given day. Only taxa with absolute log2-fold change values ≥ 0.25 were considered informative.

### qPCR analysis of ARG abundance within fecal bacterial communities

To evaluate the difference in abundance of antimicrobial resistance genes (ARGs) between and within treatment groups on days 7, 11, and 14 of the study, qPCR targets including tetracycline (*tetW, tet32*) and aminoglycoside (*aph2’-id*) antibiotic resistance genes were examined. The three gene targets were included as they were previously found at a higher abundance in swine administered therapeutic oxytet in-feed for 7 days compared to a control group (3).

qPCR assays were carried out using the QuantStudio 5 Real-Time PCR System (Thermo Fisher Scientific, San Jose, CA) and previously validated qPCR primers (3). A relative standard curve method was used for quantification. The relative standard was generated using a series of six 10-fold dilutions of fecal bacterial DNA (concentration ranging from 20 ng/μL to 2x10^-5^ ng/μL) prepared from an unpublished study that was positive for the presence of *tet32*, *tetW*, and *aph2* genes. All experimental DNA samples were run in duplicate at 4 ng per well on a 96-well PCR plate. All reactions included 10 μL iTaq Universal SYBR Green Supermix (Bio-Rad, Hercules, CA), 1 μL 10 nM forward/reverse primers, and nuclease-free water to a total volume of 20 μL per well, according to manufacturer’s recommendations. Thermal cycling conditions for *tet32, tetW*, and *aph2*-specific qPCR assays consisted of: 95°C hold for 2 min, followed by 40 cycles of 95 °C for 15 s and 60.8 °C for 60 s/cycle. Each specific target amplicon was verified by presence of a single melting temperature peak. Control reactions with no DNA template were run with each primer set to ensure the absence of non-specific primer dimers. The detection limit of the qPCR assay was set at 32 cycles for all three primer sets.

GraphPad Prism v. 8 (GraphPad Software, San Diego, CA) was used for data analysis. DNA concentrations were converted to log values using the formula: C = log(X) + 5 – log(2), where X is the DNA concentration, and C is the assigned log value. The percent coefficient of variation (CV) was calculated within each assay for each sample of each gene as: % CV = (standard deviation of A/average of A) x 100, where A is the average Ct of a set of replicates for each sample or the average C (generated by the formula above) of a set of replicates for each sample. Any sample CVs with values < 10% were considered acceptable and retained for further analysis.

To compare the abundances of the ARGs between groups among each sampling day, a mixed-effects model with Geisser-Greenhouse correction and Tukey’s multiple-comparisons test was applied to the unbalanced data using the mean log values of the samples from each group.

To compare the abundances of the ARGs between days among each group, because the data are balanced in this type of comparison, a two-way ANOVA with repeated measures model with Geisser-Greenhouse correction and Tukey’s multiple-comparisons test was applied using the mean log values of the unknowns from each group. To determine the total log10 relative abundances of ARGs for each group, the cumulative area under the log curve (AULC) was calculated for each piglet by combining days 7, 11, and 14 using the trapz function (R pracma package). Tukey’s multiple-comparisons test was used to compare the AULC between groups (*P* < 0.05).

## Data and code availability

Data and scripts are available through the USDA-FSEPRU github repository https://github.com/USDA-FSEPRU/FS9 and the sequencing data are available through the NCBI sequence read archive under BioProject PRJNA693865.

## Results

### Varying level of impact of respiratory co-infection on pig health and gut bacterial community

The study first examined the effects of co-infection of nursery-age pigs with *B. bronchiseptica* and *P. multocida* on pig health and gut bacterial community by examining samples collected on days 0, 7, 11, and 14 (Figure 1). Pigs were weighed on day 0 of the study and again at day 11 and 14, which was 4 and 7 days after initiation of oxytet treatment. At the start of the study, there were no significant differences in weight across the treatment groups (Figure 2A, Table S1). Average daily gain (ADG) was calculated for 11 and 14 days of the study. The ADG of INFnm was significantly lower than NONINFnm (*P* < 0.01) over 11 and 14 days of the trial (Figure 2B and 2C). The lung lesion scores of NONINFnm and INFnm were not significantly different on day 11 (Figure S1). However, on day 14, INFnm had significantly higher lung lesion scores (2.42 ± 2.55 [Mean ± SD]) than NONINFnm (0.42 ± 0.58) (*P* = 0.01) (Figure S1). Thus, reduced growth and greater lung lesion severity of INFnm compared to NONINFnm suggests the infection had a negative impact on the health of the animal.

To examine how co-infection of the pig host with *B. bronchiseptica* and *P. multocida* affected the GI microbiota community structure, the PERMANOVA pairwise comparisons tests (24) were performed on 16S rRNA gene amplicon data of INFnm and NONINFnm using the treatment-by-day variable. The fecal communities of INFnm to NONINFnm were compared on day 7, 11 and 14 of the study and not no significant were noted (data not shown). Thus, infection had minimal impact on the overall gut microbial community.

Though PERMANOVA pairwise comparisons results showed that, broadly-speaking, no differences were observed between the INFnm and NONINFnm pigs’ fecal microbial communities, there were individual operational taxonomic units (OTUs) more abundant in one group relative to another. To investigate how co-infection impacted the abundance of specific bacterial taxa in INFnm and NONINFnm, the fecal microbiota of both groups was analyzed at the order and genus level for each time point. Among the list of organisms that showed significant changes in abundance between the two groups (Table S2), those with log2-fold changes above 0.25 were further examined. No bacterial taxa were significantly differentially abundant between the two groups at the order level (data not shown), while at the genus level, three genera on day 7 showed significant changes in abundance. *Prevotellaceae_UCG-001* (log2-fold change = 3.01, *P* < 0.01), *Muribaculaceae_unclassified* (log2-fold change = 2.45, *P* < 0.01), and *Parabacteroides* (log2-fold change =1.66, *P* < 0.01) were relatively more abundant in INFnm than NONINFnm (Table S2). No other taxa with a log2-fold change greater than 0.25 were detected for INFnm and NONINFnm on all other days sampled in the study (days 11 and 14). These data indicated that co-infection changed the GI microbial community for a very limited amount of time.

### Oxytet administration, regardless of route, did not significantly impact colonization of B. bronchiseptica and P. multocida but improved animal health

The colonization levels of *B. bronchiseptica* and *P. multocida* from INFinject, INFfeed, and INFnm groups were compared on day 7, which was immediately prior to oxytet treatment, and days 11 and 14 (4 and 7 days post-oxytet treatment) (Table S3). Prior to antibiotic administration, colonization levels of the two challenge organisms in nasal samples were not significantly different (Table S3). Following oxytet administration, there was minimal impact on colonization levels in any tested respiratory site. Specifically, pairwise comparisons of *B. bronchiseptica* in the lung, trachea, tonsil, and nasal cavity were conducted between the three infected groups and no differences in colonization were detected. The only difference detected for *P. multocida* colonization was on day 11 in the nasal cavity where INFfeed group had significantly higher *P. multocida* levels than INFinject (*P* = 0.0054). On day 14, the lung lesion score of INFnm (Mean ± SD: 2.42 ± 2.55) group was significantly higher than that of pigs in NONINFnm (0.42 ± 0.58) (*P* ≤ 0.05) group; however, lung lesions in INFfeed (0.33 ± 0.64) and INFinject (0.39 ± 0.46) were not significantly different than lesions in the NONINFnm group (Figure S1). Thus, pigs receiving oxytet displayed minimal lung lesions comparable to the NONINFnm group.

Pigs were weighed prior to *B. bronchiseptica* inoculation (day 0) and there were no significant differences between pigs assigned to different treatment groups (Figure 2A). The average weight overall was 14.3 ± 1.78 lbs. Pigs were weighed again after oxytet administration (days 11 and 14) to assess the impact of treatment on ADG between start of study and different days post treatment. Pigs in the NONINFnm group had an ADG of 0.60 ± 0.18 lbs over 11 days and 0.72 ± 0.18 lbs over 14 days (Figure 2B and 2C), growing an average of 10.8 ± 2.84 lbs over the two-week study. Pigs in the INFnm group had an ADG of 0.37 ± 0.17 lbs over 11 days and 0.46 ± 0.14 lbs over 14 days (Figure 2B and 2C), only growing an average of 6.83 ± 2.10 lbs over the two-week study. There was a significant difference in ADG between NONINFnm and INFnm over 11 (*P* < 0.0001) and 14 (*P* = 0.006) days. The ADG of INFfeed pigs fell between tat of INFnm and NONINFnm pigs but was not significantly different from either of these groups at either timepoint (*P* > 0.09; Figure 2). On the other hand, the ADG of INFinject group was significantly higher than the ADG of INFnm group over 11 and 14 days of trial (*P* < 0.01) (Figure 2B and 2C). Pigs in the INFinject group had an ADG of 0.64 ± 0.20 lbs over 11 days and 0.72 ± 0.16 lbs over 14 days (Figure 2B and 2C), growing an average of 10.8 ± 2.54 lbs over the two-week study. Collectively, these results suggest that injected oxytet, unlike in-feed oxytet, helped to minimize the negative effects of co-infection on weight gain.

### Oxytet concentrations in the plasma, lung, and nasal cavity were dependent on administration route

Oxytet levels in the lung tissue, nasal cavity, and plasma were examined to confirm administration of antibiotic and to assess how administration route affected distribution and concentration of oxytet in the tissues sampled. On day 11 (4 days after oxytet initiation), INFfeed and INFinject groups had similar levels of oxytet in lung tissue (Figure 3A) and levels were significantly higher than INFnm (no oxytet detected) (*P* < 0.001) (Table 1). On day 14 (7 days after oxytet initiation), though INFinject had significantly lower levels of oxytet in the lung relative to INFfeed (*P* < 0.001) (Table 1, Figure 3A), but it was significantly higher than INFnm (*P* = 0.027) (Table 1). Lung oxytet levels stayed consistent over the course of in-feed treatment, as levels were not different on days 11 and 14 of the trial for INFfeed group (Mann-Whitney, *P* > 0.05). The body weight of INFfeed pigs correlated strongly with lung oxytet levels on day 11 (*P* = 0.0004, R^2^ = 0.8093) (Figure 3B), but not on day 14 despite the continued presence of oxytet in feed and consistent levels of oxytet in the lung (Figures 3A and 3B). As for INFinject, there was no significant correlation between body weight and lung oxytet level on days 11 (*P* = 0.5154, R^2^ = 0.05472) or 14 (*P =* 0.4021, R^2^ = 0.08914) (Figure 3B).

**Table 1.** Mean (+SE) oxytetracycline concentration by treatment for each tissue on day 11 and 14 of treatment as measured by LC-MS. Plasma and nasal (ng/ml) and lung (ng/2g)

|  | Day 11 |  |  | Day 14 |  |  |
| --- | --- | --- | --- | --- | --- | --- |
|  | Plasma | Lung | Nasal | Plasma | Lung | Nasal |
| INFnm | 0 ± 0 | 0 + 0 | 9 + 0 | 0 + 0 | 0 + 0 | 10 + 0 |
| INFfeed | 116.4 +<br>4.08 | 206.4 +<br>9.28 | 239.8 +<br>9.95 | 117.6 +<br>2.74 | 216.7 +<br>6.24 | 228.8 + 12 |
| INFinject | 415.4 +<br>11.99 | 203.5 +<br>6.55 | 10 + 0 | 377.5 +<br>12.98 | 45.63 +<br>1.42 | 10.72 +<br>0.15 |

In the nasal cavity, INFfeed had a significantly higher level of oxytet in the nasal wash than INFinject (*P* < 0.001) (Table 1, Figure 3C) and INFnm (Table 1). In fact, the levels of nasal oxytet detected in INFinject were below the limit of detection (10 ng/mL) on all days tested. In addition, comparison of the nasal oxytet levels in INFfeed on days 11 and 14 with the Mann-Whitney test showed the levels were similar (*P* > 0.05). This shows that nasal oxytet levels, much like in the lungs, stayed consistent over the course of in-feed treatment. No correlation was found between nasal oxytet level and body weight for pigs given oxytet in-feed on either day 11 or 14 (Figure 3D).

Plasma oxytet levels between INFfeed and INFinject were significantly different from one another on days 11 and 14 (*P* < 0.01) (Figure 3E). Oxytet administered to pigs by injection yielded the highest level of oxytet in plasma on both days 11 and 14, followed by pigs given oxytet in-feed (Table 1, Figure 3E). Plasma levels of oxytet stayed consistent from day 11 to 14 for both INFfeed and INFinject groups (Mann-Whitney, *P* > 0.05). In addition, body weight correlated with oxytet concentrations in INFinject (linear regression model, *P* = 0.003, R^2^ = 0.687) and INFfeed (*P* = 0.0452, R^2^ = 0.4127) groups on day 11 (Figure 3F). On day 14, body weights did not correlate with oxytet concentrations (INFfeed: *P* = 0.3872, R^2^ = 0.09462; INFinject: *P =* 0.0843, R^2^ = 0.3267). Similar to correlation trends observed between body weight and lung oxytet levels, the correlation between body weight and plasma oxytet levels were also short-term.

### GI bacterial communities were differentially affected by oxytet dose regimen when treating for respiratory infection

The effects of each dose regimen of oxytet on the GI microbial community when treating pigs co-infected with *B. bronchiseptica* and *P. multocida* were also examined. Differences in GI community composition between the groups were determined using PERMANOVA pairwise comparisons tests (24), focusing on the treatment effect within each day (Table 2) . On days 7 and 11, both INFinject and INFfeed showed no differences in fecal microbial community composition relative to INFnm. However, by day 14, gut communities in the INFinject fecal group differed significantly from those in the INFnm group (*P* = 0.045) (Table 2, Figure 4A). A similar magnitude of change in the INFfeed gut community relative to INFnm was observed on day 14 that approached significance (*P* = 0.084) (Table 2, Figure 4A). Thus, differences in the fecal communities of both oxytet-treated groups emerged at the end of the study.

**Table 2.** Magnitude of difference in fecal community structure (F statistic) of INFfeed and INFinject relative to INFnm on days 7, 11, and 14 of the study.

| Day | Treatment | F statistic | R<sup>2</sup> | *P*-value<sup>838</sup> |
| --- | --- | --- | --- | --- |
| 7 | INFfeed | 1.151 | 0.103 | 0.567<sup>839</sup> |
|  | INFinject | 0.881 | 0.047 | 0.57375<sup>840</sup> |
| 11 | INFfeed | 0.956 | 0.107 | 0.57375 |
|  | INFinject | 0.987 | 0.082 | 0.567 |
| 14 | INFfeed | 1.783 | 0.090 | 0.084 |
|  | INFinject | 2.208 | 0.109 | 0.045 |

To investigate how the oxytet treatment and co-infection impacted the abundance of specific bacterial taxa relative to INFnm group, counts of OTUs from fecal samples were analyzed at the order level on days 7, 11, and 14. Among the list of organisms with significant log2-fold changes in abundance between the groups (Table S4), only 6 OTUs were identified on day 14 that had absolute log2-fold changes greater than 0.25 (Figure 4B). There were decreased abundances of *Rhodospirillales* (log2-fold change = 6.21, *P* < 0.05), *Verrucomicrobiales* (log2- fold change = 4.74, *P* < 0.01), *Coriobacteriales* (log2-fold change = 1.01, *P* < 0.05), and *Gammaproteobacteria_unclassified* (log2-fold change = 0.42, *P* < 0.05) in INFfeed relative to INFnm (Figure 4B). *Verrucomicrobiales* (log2-fold change = 6.00, *P* < 0.01) and *Mollicutes_RF39* (log2-fold change = 2.13, *P* < 0.05) were relatively less abundant in INFinject than INFnm (Figure 4B). The bacterial taxa detected on day 14 may explain why there were differences in the overall fecal communities of INFinject and INFfeed relative to INFnm, as described earlier.

### Oxytet administration route impacted abundance of antimicrobial resistance genes in feces

The relative abundances of antimicrobial resistance genes (ARG) encoding aminoglycoside resistance (*aph2*) and tetracycline resistance (*tetW* and *tet32*) were examined in fecal DNA of INFnm, INFinject, and INFfeed animals on days 7, 11, and 14 to evaluate the impact of oxytet administration route on ARG abundance. When comparing between days within each treatment group, INFfeed and INFinject had significantly lower *tet32* abundances on day 11 relative to day 14 (*P* < 0.01) (Figure 5A, Table S5). As for *tetW* and *aph2* abundance, only INFinject had significantly higher abundances of the two genes on day 14 relative to days 7 and 11 (*P* < 0.05) (Figure 5B-C, Table S5). These findings suggest that the changes in *tet32* abundance within INFfeed and INFinject, and changes in *aph2* and *tetW* abundances within INFinject fluctuated through the experiment, and were at their highest relative level at the end of the trial. When comparing between the three infected groups within each sampling day, *tet32, tetW,* and *aph2* abundances were significantly reduced in INFinject group relative to INFnm on day 11 (*P* < 0.05) (Figure 5D-F, Table S5), with no difference in the INFnm between day 7 and day 11. INFinject had significantly higher *aph2* abundance relative to INFfeed on day 14 (Figure 5F, Table S5). Overall, the mean abundances of the three ARGs in all three groups increased slightly on the last day of the study.

We also compared the total relative abundances of the three ARGs for each group by combining log10 relative abundances of each ARG from days 7, 11, and 14 and calculating the area under the log curve (AULC) (Figure 6). No significant difference in AULC measurements were noted among the three infected groups for *tet32* and *aph2* (Figure 6A and C). However, the AULC measurement of INFinject was significantly lower than INFfeed and INFnm (*P* < 0.05) for *tetW* (Figure 6B). INFinject in general had the lowest mean AULC values relative to the other two groups for all three ARGs.

## Discussion

Antibiotic stewardship efforts include selecting an appropriate administration route for an antibiotic with effective therapeutic response but also limited risk of antimicrobial resistance emergence; however, these and additional stewardship efforts remain a scientific challenge (27). Oxytetracycline (oxytet) can be administered through multiple routes to pigs. It belongs to the tetracycline class of antibiotics, a class commonly used for prophylactic and therapeutic treatment in growing pigs in the U.S. in 2016 and 2017 (28). Oxytet is labeled for treatment of a number of diseases including bacterial enteritis caused by *Escherichia coli* and bacterial pneumonia caused by *Pasteurella multocida*.

A recent study investigated how the route of oxytet administration may differentially impact the GI microbiome when administered to weaned piglets (3). Both injected and infeed oxytet had significant effects on the gut microbiota. In addition, in-feed oxytet had greater effect on the overall gut bacterial community, it changed the relative abundance of multiple bacterial taxa, and increased the prevalence of specific antimicrobial resistance genes. However, the animals in the previously published study were treated prophylactically. It is unknown whether each route of oxytet is equally effective at treating an animal with an infection, and what additional effects each route has on a diseased animal’s health and GI microbiota. This study aimed to address those questions by examining the impact of oxytet administration route on pigs challenged with respiratory pathogens *B. bronchiseptica* and *P. multocida*. We first evaluated how infection alone affected the health and gut microbiota of weaned pigs. Second, we evaluated how effective each oxytet route was at clearing the infection and also how impactful treatment and infection were on the following: oxytet levels at various sites of the pig, health of the pig by lung lesion severity and growth rate, and lastly, pig gut microbial community profile using 16S rRNA sequencing targeting the V4 region.

A small number of reports showed respiratory infection can have significant impact on the pig gut microbiota (29–31), though these studies only examined infection caused by porcine reproductive and respiratory syndrome virus (PRRSV), which can enter through the respiratory tract with subsequent viremia. One reported the virulence of a PRRSV strain can influence disease severity and abundance of commensal organisms in the pig gut microbiome (32). There was significant positive correlation between the depletion of desirable commensals and severe infection caused by the virulent PRRSV strain compared to the less virulent strain (32). Like the less virulent PRRSV strain, the *B. bronchiseptica* and *P. multocida* strains used in our study caused mild clinical signs in pigs as a result of their low virulence potential (S. Brockmeier, personal correspondence). We also observed minimal differences in the overall fecal community profile and the number of differentially abundant bacterial taxa between infected and non-infected pigs. The minimal effects of respiratory infection on the pig GI microbial communities may be attributed to the limited pathogenesis of our two strains. These nominal changes in our infected pig group were also not entirely unique. In a study conducted in foals, the authors observed minimal impact of respiratory infection on the GI microbiota (33). Foals were infected with a respiratory pathogen, *Rhodococcus equi*, and regardless of whether the foals displayed no clinical signs, mild clinical signs, or severe clinical signs, no significant changes in the fecal microbial composition or diversity were detected (33).

Whilst co-infection with *B. bronchiseptica* and *P. multocida* caused moderate impact on the pigs’ health, there was little to no reduction in colonization levels of either organism measured from various tissue sites of the pigs after initiation of oxytet treatment. One reason may be the bacteriostatic function of oxytet (34). A second reason may be the poor bioavailability of oxytet when given in feed. Consistent with our previous study (3), the INFfeed pigs had lower plasma oxytet levels than INFinject pigs. The slow disintegration of pellets and release of oxytet from the pellets may affect oxytet absorption (35). Food components may affect its availability, thus the clinically effective dose of in-feed oxytet against *P. multocida* is not achieved (35). A third reason could be the oxytet levels detected in plasma, nasal, and lung measured all throughout the current experiment (Table 1) were below the established minimum inhibitory concentration (MIC) of 0.5µg/mL for *B. bronchiseptica* and *P. multocida* (data not shown), though we did not measure before day 4 after oxytet treatment (levels may have been above MIC at earlier timepoint). The formulations used in this study were not effective in reducing colonization. Two recent studies found that the depot formulation of injected oxytet licensed to treat porcine respiratory diseases at the recommended single dose of 20 mg/kg would not attain levels necessary for direct killing of *P. multocida* (36, 37). The dosage we used for injected and in-feed oxytet were equivalent to 20 mg/kg and 22 mg/kg, respectively.

Although the direct killing action of oxytet may not always be achieved with manufacturer’s recommended dosage, tetracyclines have other actions that may contribute to its efficacy, such as altered production of virulence factors (36, 37). In our study, even though the oxytet levels did not reduce colonization levels of the respiratory organisms, there were potential actions of efficacy from oxytet treatment including reduced lung lesion severity by both routes and minimized negative effects of co-infection on weight gain by injected oxytet. A similar observation was seen in pigs treated for a respiratory infection with injected oxytet (38). Pigs were prophylactically treated with a 20 mg/kg dose of long-acting intramuscular injection of oxytet (similar to the dose used in this study) and then challenged intranasally with *Actinobacillus pleuropneumoniae* either 24 h or 48 h post-treatment (38). The injected oxytet dose significantly reduced mortality and produced less severe clinical signs, such as lung lesions, compared to infected control pigs; however, *A. pleuropneumoniae* was not enumerated so it’s unclear if the effect was due to bacterial burden or limiting virulence.

Changes in the fecal communities of both oxytet-treated groups relative to INFnm were limited. Injected oxytet had a significant but small effect on the fecal microbiota towards the end of the study while in-feed oxytet showed no significant changes in its fecal microbiota relative to INFnm. Ricker et al., however, saw significant impact on the fecal microbiome by both routes of oxytet relative to the non-medicated group post-oxytet treatment (3). Relative to the non-medicated group, the in-feed oxytet group also had much larger fecal microbiome differences than injected oxytet (3). In our study, the magnitude of change in the INFfeed group’s fecal microbiota (Table 2, days 11 and 14) was not only lower than INFinject, it was also much lower than what we previously observed in our in-feed oxytet group. On the other hand, injected oxytet treatment, given prophylactically (3) or therapeutically in this study, had limited impact on the fecal microbiota community. As noted in our prior work, we anticipate the limited impact of injected oxytet on the microbiota was due to the limited amount of oxytet in the gut lumen (measured as fecal oxytet levels) (3).

As for the relative abundances of *tet32, tetW,* and *aph2* resistance genes in the infected groups, INFinject had significantly lower abundances of all three ARGs compared to INFnm 4 days post-oxytet administration (day 11). However, this difference was not sustained and no longer observed 7 days post-oxytet treatment (day 14). The abundances of the three ARGs in INFfeed remained unchanged relative to INFnm over the course of oxytet treatment. Ricker et al. observed significantly greater abundances of *tetW* and *aph2* in their in-feed oxytet group compared to the non-medicated group 7 days after administering oxytet in-feed. A similar study also observed greater abundance of tetracycline-resistance genes in fecal samples from weaned pigs following 7-day prophylactic in-feed oxytet treatment (39). It’s possible the respiratory infection may have slightly reduced feed intake when compared to prior, prophylactic studies and thus, altered the impact on microbiota and relative ARG abundance. Support for this comes from a lack of significant difference in weight between the INFfeed to INFnm or NONINFnm group, but it is worth noting oxytet was detected in lung, nasal wash and plasma of the INFfeed group, suggesting the antibiotic was absorbed. Injected oxytet may lead to a pronounced change in abundance of organisms carrying tetracycline- or aminoglycoside-resistance relative to INFnm and possibly contribute to the significant changes in overall fecal community structure observed on day 14. Thus far, no studies have examined the differential effects of antibiotic administration route on the fecal microbial community and resistance gene abundance of infected animals. Additional studies are needed to better understand the relationship between microbiota structure and ARG abundance.

Overall, injected oxytet resulted in slightly better outcomes than in-feed oxytet in this study, such as decreased abundances of select ARGs (Figures 5 and 6) and protection of infected pigs from worsened disease. While injected oxytet may seem more preferable to in-feed oxytet for treating pigs with a respiratory infection, there were several limitations to the study. Due to limited availability of animal resources, we were unable to include a group of pigs that were non-infected and treated with only oxytet by either injected or in-feed route. An inclusion of such a group would confirm whether some of the differences observed were mainly due to the specific oxytet treatment alone or in combination with other factors. The piglets used in this study likely received maternal antibodies against *B. bronchiseptica* prior to receipt, given it’s ubiquitous nature in commercial pigs, and possibly *P. multocida* antibodies as well. This may explain why subsequent challenge of pigs with *P. multocida* did not elicit a severe respiratory infection. The colonization levels of both respiratory pathogens prior to oxytet treatment or in INFnm group were also relatively low compared to previous studies that have used this infection model (17, 40). This study did not measure fecal oxytet concentration, and thus we cannot resolve whether the microbiota shifts observed in the study were also related to the concentration of oxytet in the gut region or a combination of factors. Future efforts should address these limitations to further clarify the efficacy of each route of oxytet treatment on infection and on the health and GI microbiome profile of the animal.

The present study observed INFnm had limited disease outcomes and minimal changes in the fecal microbiota compared to NONINFnm. In addition, administration of oxytet by either route was not efficacious in reducing pathogen burden in infected animals. However, both oxytet routes had some measurable positive impact on animal health and showed limited impact on the fecal microbial community and relative abundance of specific tetracycline genes. Even so, one must weigh between the various drug options for their treatment efficacy, their effects on the animal’s performance, and their impact on gut microbiota including the prevalence of antimicrobial resistance and opportunistic pathogens. The results from this study contribute to continuing efforts to identify antimicrobial applications that can minimize impact on gut microbiome without compromising treatment efficacy in food-producing animals.

## Ethics and Conflict of Interest Statement

The animal study was reviewed and approved by USDA-National Animal Disease Center Animal Care and Use Committee. Mention of trade names or commercial products in this publication is solely for the purpose of providing specific information and does not imply recommendation or endorsement by the U.S Department of Agriculture. USDA is an equal opportunity provider and employer.

## Dedication

This article is dedicated to the memory of Heather K. Allen.

## Acknowledgements

We are deeply grateful for the animal caretakers at the NADC, and Zahra Bond, Jennifer Jones, Steve Kellner, Amber Miranda, Morgan Smith, and Elli Whalen for assistance in collecting and/or preparing samples for sequencing, for David Alt for sequencing the samples, and for investigators from the Food Safety and Enteric Pathogens Research Unit for fruitful discussions in data interpretation.

## Figure Captions

Figure 1. Diagram of experimental timeline. The table describes the four groups from the study, the infection status, and/or oxytetracycline (oxytet) treatment administered.

Figure 2. Comparison of pig weight (pounds) on day 0 (A), average daily gain (pounds) between INFfeed, INFinject, INFnm, and NONINFnm on day 11 (B), and day 14 (C). *P* < 0.05 indicates significant difference (*).

Figure 3. Oxytet levels detected in lung, nasal, and plasma of INFinject and INFfeed pigs sampled on days 11 and 14 (A, C, E) and relative to body weight (lbs) (B, D, F). Tissue samples and weight (lbs) were collected at the indicated time point (days 11 and 14) and LC-MS was performed to determine oxytet concentrations (y-axis) in tissues. The INFinject group received a single therapeutic dose on day 7 and the INFfeed group received a therapeutic dose in-feed up from day 7 to day 14 as described in the methods.

Figure 4. Impact of oxytet administration route and infection on fecal microbial community structure on days 7, 11, and 14 (A), and the identification of significantly less abundant bacterial taxa at the Order level (*P* < 0.05) in the fecal microbiota of INFfeed and INFinject relative to INFnm on day 14 (B). NMDS ordinations of fecal 16S rRNA gene-based Bray-Curtis dissimilarities were calculated from rarerified OTU abundance data (clustered at 97% similarity). Gray-colored points represent pig samples from other days. Ordination stress = 0.182. Comparisons in (B) are relative to INFnm group.

Figure 5. Impact of oxytet administration route on relative abundance of antibiotic resistance genes (ARG) tet32 (A, D), tetW (B, E), and aph2 (C, F) in pig feces. Comparisons of mean log10 relative abundance of each ARG (y-axis) were made between days for each treatment group (A, B, C) or between treatment groups for each day (D, E, F). Supplemental Table 5 includes all pairwise comparison results.

Figure 6. Comparison of total area under the log curve (AULC) of log10 ARG relative abundances of tet32 (A), tetW (B), or aph2 (C) over sampling days 7, 11, and 14 for INFfeed, INFinject, and INFnm. *P* < 0.05 indicates significant difference (*).

Figure S1. Comparison of mean scores for lung lesion severity on days 11 and 14 due to co- infection with *B. Bordetella* and *P. multocida* and oxytetracycline administration. The x-axis displays the days treatment groups were sampled and y-axis shows weighted average score of lung lesion, with 0 = no lung lesions detected. Dots represent individual pig samples. *P* < 0.05 indicates significance (*).

## Supplemental Tables

Table S1. Metadata of animals used in study, weight, and average daily gain.

Table S2. List of OTU with log2-fold changes in abundance greater than 0.25 between NONINFnm and INFnm.

Table S3. Colonization levels of *Bordetella bronchiseptica* or *Pasteurella multocida* in various tissues detected on days 11 and 14.

Table S4. List of OTU with log2-fold changes in abundance greater than 0.25 for INFfeed and INFinject relative to INFnm.

Table S5. Pairwise comparisons of mean log10 relative abundances of *tet32, tetW, aph2* genes in INFfeed, INFinject, and INFnm groups on days 7, 11, and 14.

---

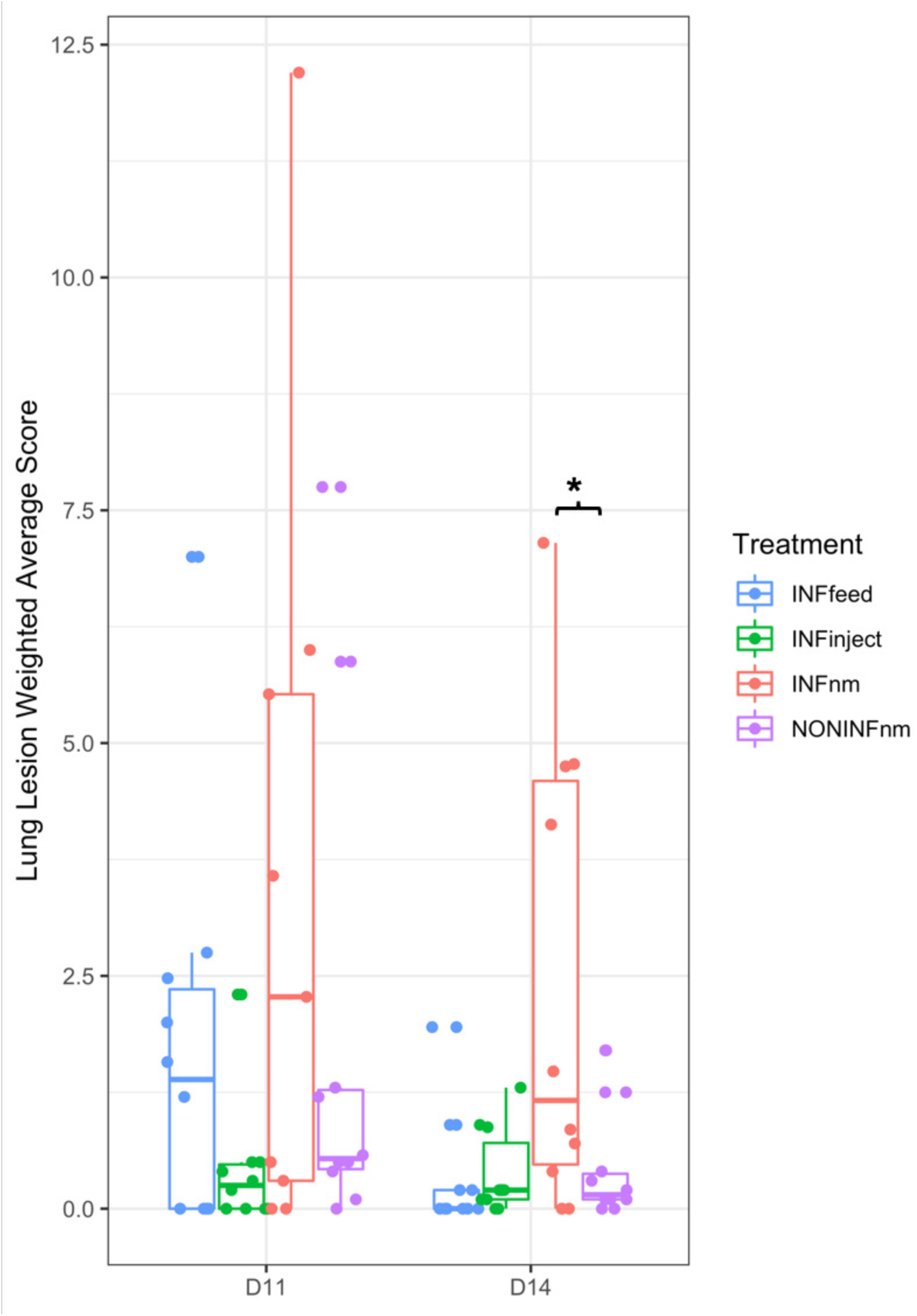

## References

1. McEwen SA, Collignon PJ. 2018. Antimicrobial Resistance: a One Health Perspective. Microbiology Spectrum 6.

2. FDA. 2015. Veterinary Feed Directive; Final Rule. Services DoHaH, https://www.federalregister.gov/documents/2015/06/03/2015-13393/veterinary-feed-directive.

3. Ricker N, Trachsel J, Colgan P, Jones J, Choi J, Lee J, Coetzee JF, Howe A, Brockmeier SL, Loving CL, Allen HK. 2020. Toward Antibiotic Stewardship: Route of Antibiotic Administration Impacts the Microbiota and Resistance Gene Diversity in Swine Feces. Front Vet Sci 7:255.

4. Zeineldin M, Aldridge B, Lowe J. 2019. Antimicrobial Effects on Swine Gastrointestinal Microbiota and Their Accompanying Antibiotic Resistome. Frontiers in Microbiology 10.

5. Holman DB, Chénier MR. 2015. Antimicrobial use in swine production and its effect on the swine gut microbiota and antimicrobial resistance. Canadian Journal of Microbiology 61:785–798.

6. Looft T, Allen HK. 2012. Collateral effects of antibiotics on mammalian gut microbiomes. Gut Microbes 3:463–467.

7. Bibbal D, Dupouy V, Ferré JP, Toutain PL, Fayet O, Prère MF, Bousquet-Mélou A. 2007. Impact of Three Ampicillin Dosage Regimens on Selection of Ampicillin Resistance in *Enterobacteriaceae* and Excretion of *bla*TEM Genes in Swine Feces. Applied and Environmental Microbiology 73:4785–4790.

8. Starke IC, Pieper R, Neumann K, Zentek J, Vahjen W. 2014. The impact of high dietary zinc oxide on the development of the intestinal microbiota in weaned piglets. FEMS Microbiology Ecology 87:416–427.

9. Luppi A. 2017. Swine enteric colibacillosis: diagnosis, therapy and antimicrobial resistance. Porcine Health Management 3:16.

10. Rhouma M, Fairbrother JM, Beaudry F, Letellier A. 2017. Post weaning diarrhea in pigs: risk factors and non-colistin-based control strategies. Acta Veterinaria Scandinavica 59:31.

11. Jensen ML, Thymann T, Cilieborg MS, Lykke M, Mølbak L, Jensen BB, Schmidt M, Kelly D, Mulder I, Burrin DG, Sangild PT. 2014. Antibiotics modulate intestinal immunity and prevent necrotizing enterocolitis in preterm neonatal piglets. American Journal of Physiology-Gastrointestinal and Liver Physiology 306:G59–G71.

12. Niederwerder MC. 2017. Role of the microbiome in swine respiratory disease. Veterinary Microbiology doi:http://dx.doi.org/10.1016/j.vetmic.2017.02.017.

13. Carlson MS, Fangman TJ. 2000. Antibiotics and Other Additives for Swine: Food Safety Considerations, on University of Missouri Extension. https://extension2.missouri.edu/g2353. Accessed April 22, 2018.

14. Granados-Chinchilla F, Rodríguez C. 2017. Tetracyclines in Food and Feedingstuffs: From Regulation to Analytical Methods, Bacterial Resistance, and Environmental and Health Implications. Journal of Analytical Methods in Chemistry 2017:1315497.

15. Brockmeier SL, Palmer MV, Bolin SR, Rimler RB. 2001. Effects of intranasal inoculation with Bordetella bronchiseptica, porcine reproductive and respiratory syndrome virus, or a combination of both organisms on subsequent infection with Pasteurella multocida in pigs. American Journal of Veterinary Research 62:521–525.

16. Nicholson TL, Shore SM, Bayles DO, Register KB, Kingsley RA. 2014. Draft Genome Sequence of the Bordetella bronchiseptica Swine Isolate KM22. Genome Announc 2.

17. Brockmeier SL, Register KB. 2007. Expression of the dermonecrotic toxin by Bordetella bronchiseptica is not necessary for predisposing to infection with toxigenic Pasteurella multocida. Veterinary Microbiology 125:284–289.

18. Loving CL, Brockmeier SL, Vincent AL, Palmer MV, Sacco RE, Nicholson TL. 2010. Influenza virus coinfection with Bordetella bronchiseptica enhances bacterial colonization and host responses exacerbating pulmonary lesions. Microbial Pathogenesis 49:237–245.

19. Hughes HR, Brockmeier SL, Loving CL. 2018. Bordetella bronchiseptica Colonization Limits Efficacy, but Not Immunogenicity, of Live-Attenuated Influenza Virus Vaccine and Enhances Pathogenesis After Influenza Challenge. Frontiers in Immunology 9.

20. Trachsel J, Briggs C, Gabler NK, Allen HK, Loving CL. 2019. Dietary Resistant Potato Starch Alters Intestinal Microbial Communities and Their Metabolites, and Markers of Immune Regulation and Barrier Function in Swine. Front Immunol 10:1381.

21. Halbur PG, Paul PS, Frey ML, Landgraf J, Eernisse K, Meng XJ, Lum MA, Andrews JJ, Rathje JA. 1995. Comparison of the pathogenicity of two US porcine reproductive and respiratory syndrome virus isolates with that of the Lelystad virus. Vet Pathol 32:648–60.

22. Kozich JJ, Westcott SL, Baxter NT, Highlander SK, Schloss PD. 2013. Development of a Dual-Index Sequencing Strategy and Curation Pipeline for Analyzing Amplicon Sequence Data on the MiSeq Illumina Sequencing Platform. Applied and Environmental Microbiology 79:5112–5120.

23. Mou KT, Allen HK, Alt DP, Trachsel J, Hau SJ, Coetzee JF, Holman DB, Kellner S, Loving CL, Brockmeier SL. 2019. Shifts in the nasal microbiota of swine in response to different dosing regimens of oxytetracycline administration. Veterinary Microbiology 237:108386.

24. Martinez Arbizu P. 2019. pairwiseAdonis: Pairwise multilevel comparison using adonis. R package version 0.3. https://github.com/pmartinezarbizu/pairwiseAdonis Accessed

25. Love MI, Huber W, Anders S. 2014. Moderated estimation of fold change and dispersion for RNA-seq data with DESeq2. Genome Biology 15:550.

26. Zhu A, Ibrahim JG, Love MI. 2019. Heavy-tailed prior distributions for sequence count data: removing the noise and preserving large differences. Bioinformatics 35:2084–2092.

27. van Doorn D. Oct. 14, 2015 2015. Why relying on antibiotics alone is not enough. Pig Progress.

28. Davies PR, Singer RS. 2020. Antimicrobial use in wean to market pigs in the United States assessed via voluntary sharing of proprietary data. Zoonoses and Public Health 67:6–21.

29. Jiang N, Liu H, Wang P, Huang J, Han H, Wang Q. 2019. Illumina MiSeq Sequencing Investigation of Microbiota in Bronchoalveolar Lavage Fluid and Cecum of the Swine Infected with PRRSV. Current Microbiology 76:222–230.

30. Niederwerder MC, Jaing CJ, Thissen JB, Cino-Ozuna AG, McLoughlin KS, Rowland RRR. 2016. Microbiome associations in pigs with the best and worst clinical outcomes following co-infection with porcine reproductive and respiratory syndrome virus (PRRSV) and porcine circovirus type 2 (PCV2). Veterinary Microbiology 188:1–11.

31. Ober RA, Thissen JB, Jaing CJ, Cino-Ozuna AG, Rowland RRR, Niederwerder MC. 2017. Increased microbiome diversity at the time of infection is associated with improved growth rates of pigs after co-infection with porcine reproductive and respiratory syndrome virus (PRRSV) and porcine circovirus type 2 (PCV2). Veterinary Microbiology 208:203–211.

32. Argüello H, Rodríguez-Gómez IM, Sánchez-Carvajal JM, Pallares FJ, Díaz I, Cabrera- Rubio R, Crispie F, Cotter PD, Mateu E, Martín-Valls G, Carrasco L, Gómez-Laguna J. 2021. Porcine reproductive and respiratory syndrome virus impacts on gut microbiome in a strain virulence-dependent fashion. Microbial Biotechnology n/a.

33. Whitfield-Cargile CM, Cohen ND, Suchodolski J, Chaffin MK, McQueen CM, Arnold CE, Dowd SE, Blodgett GP. 2015. Composition and Diversity of the Fecal Microbiome and Inferred Fecal Metagenome Does Not Predict Subsequent Pneumonia Caused by Rhodococcus equi in Foals. PLOS ONE 10:e0136586.

34. Loree J LS. 2021. Bacteriostatic Antibiotics. *In* (ed), StatPearls StatPearls Publishing, Treasure Island, FL. https://www.ncbi.nlm.nih.gov/books/NBK547678/.

35. Mevius DJ, Vellenga L, Breukink HJ, Nouws JFM, Vree TB, Driessens F. 1986. Pharmacokinetics and renal clearance of oxytetracycline in piglets following intravenous and oral administration. Veterinary Quarterly 8:274–284.

36. Dorey L, Hobson S, Lees P. 2017. What is the true in vitro potency of oxytetracycline for the pig pneumonia pathogens Actinobacillus pleuropneumoniae and Pasteurella multocida? Journal of Veterinary Pharmacology and Therapeutics 40:517–529.

37. Dorey L, Pelligand L, Cheng Z, Lees P. 2017. Pharmacokinetic/pharmacodynamic integration and modelling of oxytetracycline for the porcine pneumonia pathogens Actinobacillus pleuropneumoniae and Pasteurella multocida. Journal of Veterinary Pharmacology and Therapeutics 40:505–516.

38. Kiorpes AL, Bäckström LR, Collins MT, Kruse GO. 1989. Comparison of conventional and long-acting oxytetracyclines in prevention of induced Actinobacillus (Haemophilus) pleuropneumoniae infection of growing swine. Canadian journal of veterinary research = Revue canadienne de recherche veterinaire 53:400–404.

39. Ghanbari M, Klose V, Crispie F, Cotter PD. 2019. The dynamics of the antibiotic resistome in the feces of freshly weaned pigs following therapeutic administration of oxytetracycline. Scientific Reports 9:4062.

40. Chanter N, Magyar T, Rutter JM. 1989. Interactions between Bordetella bronchiseptica and toxigenic Pasteurella multocida in atrophic rhinitis of pigs. Research in Veterinary Science 47:48–53.

